# Spatial phylogenetics of Fagales: Investigating the history of temperate forests

**DOI:** 10.1101/2023.04.17.537249

**Authors:** R.A. Folk, C.M. Siniscalchi, J. Doby, H.R. Kates, S.R. Manchester, P.S. Soltis, D.E. Soltis, R.P. Guralnick, M. Belitz

## Abstract

**Aim:** Quantifying the phylogenetic diversity of temperate trees is essential for understanding what processes are implicated in shaping the modern distribution of temperate broadleaf forest and other major forest biomes. Here we focus on Fagales, an iconic member of forests worldwide, to uncover global diversity and endemism patterns and investigate potential drivers responsible for the spatial distribution of fagalean forest communities.

**Location:** Global.

**Taxon:** Fagales.

**Methods:** We combined phylogenetic data covering 60.2% of living species, fine-scale distribution models covering 90% of species, and nodulation data covering all species to investigate the distribution of species richness at fine spatial scales and compare this to relative phylogenetic diversity (RPD) and phylogenetic endemism. Further, we quantify phylogenetic betadiversity and bioregionalization of Fagales and determine hotspots of Fagales species engaging in root nodule symbiosis (RNS) with nitrogen-fixing actinomycetes.

**Results:** We find the highest richness in temperate east Asia, eastern North America, and equatorial montane regions of Asia and Central America. By contrast, RPD is highest at higher latitudes, where RNS also predominates. We found a strong spatial structuring of regionalizations of Fagales floras as defined by phylogeny and traits related to RNS, reflecting distinct Northern and Southern Hemisphere floras (with the exception of a unique Afro-Boreal region) and highly distinct tropical montane communities.

**Main conclusions:** Species richness and phylogenetic regionalization accord well with traditional biogeographic concepts for temperate forests, but RPD does not. This may reflect ecological filtering specific to Fagales, as RNS strategies are almost universal in the highest RPD regions. Our results highlight the importance of global-scale, clade-specific spatial phylogenetics and its utility for understanding the history behind temperate forest diversity.

## INTRODUCTION

The distribution of today’s forest biomes is profoundly shaped by the division of tropical and extratropical floras into distinct phytogeographical domains, with limited mutual migration imposed by niche conservatism (Donoghue, 2008; Folk et al., 2020; Takhtajan et al., 1986; Wiens & Donoghue, 2004) leading to distinctive macroevolutionary histories in these regions (Axelrod, 1966; Economo et al., 2018; Edwards et al., 2017; Schubert et al., 2019; Sun et al., 2020; Wolfe, 1987). Angiosperm tree species richness outside the tropics can generally be summarized as a distribution in isolated regions along a belt in the Northern Hemisphere, reaching its highest diversity in eastern Asia, its second-highest diversity in eastern North America, and minor diverse areas stretching from Asia westward to the Irano-Turanian region, the Caucasus, and Anatolia in Eurasia and from eastern North America to the California Floristic Province. The temperate areas of the Southern Hemisphere able to support angiosperm forests are more limited in spatial extent, with the primary diversity areas comprising southern South America, Tasmania, and New Zealand, primarily at higher elevations. Although physiological challenges such as frost and elevated seasonality are important for governing the distribution of temperate floras, the diversity of tree species observed in today’s temperate forests is more unevenly distributed than would be suggested by these factors alone (see Fig. 2 in Folk et al., 2020).

The discontinuous and imbalanced distribution of temperate forests has been hypothesized to reflect refugial areas remaining after the disappearance of ancient polar forests (Engler, 1905; Axelrod, 1983; Manchester, 1999; Wen, 1999; Wolfe, 1975). Under this hypothesis, hotspots of temperate forest diversity would represent climatically buffered, topographically complex areas that promote diversification while lessening the impact of extinction. Persistence of the original lineage composition of an ancient northern temperate tree flora is considered greatest in eastern Asia, while aridification and glaciation resulted in the loss of many lineages elsewhere and a near-total loss in western Europe and western North America. While refugial areas of ancient temperate forests have long been hypothesized to explain centers of angiosperm tree diversity (Manchester, 1999; Wen, 1999; Wolfe, 1975), as informed by changes in distribution patterns documented in the fossil record, alternative hypotheses, such as ancestral origins in current diversity hotspots or ecological filtering (Cavender-Bares et al., 2004), are also plausible and largely untested.

Critical to testing refugial or other hypotheses of how centers of temperate forest diversity are maintained is mapping areas of tree diversity and connecting these data to phylogenetic hypotheses (Allen et al., 2019; Mishler et al., 2020; Scherson et al., 2017; Thornhill et al., 2017). Different measures of biodiversity, including not only species richness but phylogenetic facets, help reveal additional aspects of lineage diversity relevant to distinguishing among hypotheses (Doby et al., 2022; D. Li et al., 2020). Using spatial phylogenetic tools (Mishler et al., 2014) to distinguish areas of endemism promises to provide additional insight into areas that harbor ancient diversity (i.e., paleoendemism, which could indicate areas of reduced extinction), or that disproportionately contain recently evolved narrow endemics (i.e., neoendemism, which could be the result of ecological filtering or recent in situ diversification). Joining diversity estimates with contemporary environmental data and with ecologically significant traits yields additional power to distinguish ecological filtering from alternative explanations. Combining measures of biodiversity with measures of external environment helps to potentially distinguish among hypotheses for the maintenance of biodiversity (Suissa et al., 2021; Thornhill et al., 2017).

Hardly a better clade could be chosen to understand the origin of forests in extratropical areas than Fagales, an iconic member of temperate forests worldwide (recognized as an order by APG [2016]). Its members include some of the most familiar Northern Hemisphere trees such as alder (*Alnus*), beech (*Fagus*), birch (*Betula*), hickory (*Carya*), oak (*Quercus*), and walnut (*Juglans*), and important plants of the Southern Hemisphere such as southern beech (*Nothofagus*) and she-oak (Casuarinaceae). Standing out among major woody clades for its ecological diversity, Fagales are broadly present across major temperate to boreal forest biomes with a significant additional presence in tropical areas, both hot lowland areas (Casuarinaceae) and cooler upland regions (especially Juglandaceae and Ticodendraceae) (Wheeler et al., 2022).

Areas of diversity for Fagales cover the major centers of diversity for temperate forests, with substantial richness in eastern Asia and eastern temperate North America, strongholds of diversity for many of the constituent families. The geographic tropics harbor lower fagalean species diversity than other regions, but contain many phylogenetically distinct fagalean lineages (e.g., *Alfaroa, Engelhardia, Trigonobalanus*, Ticodendraceae) as well as additional centers of diversity for otherwise temperate genera (Axelrod, 1983), particularly in montane Central America and Malaysia. Fagales are similarly important within the limited extent of temperate forest in the Southern Hemisphere, with Nothofagaceae occurring across southern South America to New Zealand and Papua New Guinea, forming an iconic Antarctic disjunction (Cook & Crisp, 2005; Hinojosa et al., 2016), and Myricaceae and Casuarinaceae covering the remaining southerly latitudes in Africa and Australasia, respectively.

Fagales have the additional feature of distinctive morphological diversity, with some groups such as Casuarinaceae difficult to relate to other clades in terms of structural homology. A standout adaptation is the presence of root nodules in some Fagales, i.e., specialized root structures that house symbiotic diazotrophic bacteria. Fagales contain three of the eight families (viz., Betulaceae, Casuarinaceae, and Myricaceae; Pawlowski & Bisseling, 1996; Pawlowski & Sprent, 2008) that nodulate, representing three of nine independent origins (Kates et al., 2022), of “actinorhizal” plants, those whose root nodule symbiosis (RNS) involves Actinomycetota (Actinobacteria) rather than the Alpha-or Betaproteobacteria (“rhizobia”) found predominantly in the legumes (Fabaceae). Recently, differences in latitudinal diversity have been identified between plants with differing bacterial partners; compared to other forms of diazotrophic symbiosis, actinorhizal RNS is most prevalent in temperate to boreal environments (Tamme et al., 2021) while other forms of diazotrophic symbiosis achieve their greatest prevalence in the tropics and subtropics. While the distinctive habitats of actinorhizal plants have been noted (Folk et al., 2020; Menge et al., 2019; Tamme et al., 2021), we do not understand the specific climatic factors that may be causative of differing latitudinal patterns among symbiosis types; clarifying these factors would shed light on the interaction between RNS and the abiotic environment.

The long history of study and strong baseline of morphological and phylogenetic information in Fagales (reviewed in Wheeler et al., 2022), as well as their presence across such a wide range of areas and habitats, provide a basis for understanding the assembly of modern temperate forest biomes and their relation to the tropics (Folk et al., 2020; Spriggs et al., 2015). While diversity mapping has seen intense interest in trees generally (e.g., Lyu et al., 2022; Segovia et al., 2020) and in focused groups of Fagales, such as oaks (Cavender-Bares et al., 2004), Fagales themselves have never been the subject of a focused spatial phylogenetic analysis. Yet understanding the distribution of phylogenetic diversity in globally distributed model clades (Cavender-Bares, 2019) is essential to determining whether results at broad phylogenetic scales generalize to different levels of comparison (Graham et al., 2018) while providing a more direct means to uncover processes that operate at shallower scales (Cavender-Bares, 2019). In Fagales, in particular, broader comparisons are critical for understanding how key traits such as RNS impacted global diversity.

Here, we assemble a view of fagalean phylodiversity that is fine-grained yet global in extent, including measures of endemism and phylogenetic beta-diversity, and relate these results to environmental factors that promote diversity and endemism. To do so, we developed a new species distribution modeling pipeline that is semi-automated and uses best practices for modeling in order to produce a robust map of fagalean diversity. Finally, we investigated and mapped the proportion of lineages within Fagales that engage in RNS, asking whether this ecological strategy may be involved in ecological filtering of phylogenetic diversity. We address four primary goals with these assembled data resources in a modeling framework: (1) assess areas and environmental conditions associated with high species richness and phylogenetic diversity; (2) distinguish patterns of neoendemism and paleoendemism; and (3) use phylogenetic regionalization analyses based on phylogenetic beta-diversity to test longstanding hypotheses of the regionalization of forest diversity. Finally, to understand the spatial distribution of a key functional trait in Fagales, we (4) identify environments that support a higher diversity of species that engage in actinorhizal RNS.

## METHODS

### Phylogenetic framework

Fagales are well-studied phylogenetically with numerous approaches to date, from Sanger sequencing (R. Li et al., 2004; Manos et al., 2007) to genomic data (Yang et al., 2021) and fossils (Larson-Johnson, 2016; Siniscalchi et al., 2023). For the purpose of spatial phylogenetics, maximizing species presence is most important (D. Li et al., 2019), so we elected to use a recent approach combining genomic data with all Fagales sequences available on GenBank (Siniscalchi et al., 2022). This tree, based on 20 regions from the nucleus, chloroplast, and mitochondrion, comprises 707 species of Fagales (60.2% of species-level diversity; Stevens, 2001 onwards) with complete genus-level representation.

Briefly, the tree was reconstructed by aggregating all GenBank data available for these 20 markers, as well as extracting these same markers from a recent large-scale sequencing initiative (Kates et al., 2022). Tree inference, performed in RAxML-NG (Kozlov et al., 2019), was constrained by a high-quality backbone based on nuclear phylogenomic data derived from Kates et al. (2022); analytical details are available in Siniscalchi et al. (2022).

### Occurrence records

Occurrence records of all Fagales found on the biodiversity discovery platforms GBIF (*GBIF*, 2020) and iDigBio were aggregated on May 21, 2020, compiling a dataset of 7,623,848 records. Next, we harmonized synonyms to a standardized species list (the NitFix names database, as reported in Folk et al., 2021) to deal with taxonomic changes that may not be up to date in GBIF or iDigBio repositories. Duplicate records and specimens, here meaning records or specimens collected at the same location, by the same collector, on the same day, were filtered to only retain single specimen records. Additionally, records were removed if they did not have coordinate values. Each species that had at least five occurrence records underwent a coordinate cleaning procedure using the *CoordinateCleaner* package (Zizka et al., 2019). During this procedure, records were removed if they: 1) had equal latitude and longitude coordinates, 2) were within 500 m of the centroids of political countries or provinces, 3) were within 0.5 degree radius of the GBIF headquarters, 4) were within 100 m of zoos, botanical gardens, herbaria, and museums based on a global database of ∼10,000 such biodiversity institutions (see Zizka et al., 2019), 5) had precisely zero values for latitude or longitude (and therefore would likely be erroneous), or 6) were greater than 1000 km away from all other records of a species. Dot maps of occurrence records were then plotted, and based on manual inspection of species occurrence maps, species that retained obvious errors underwent manual occurrence record filtering.

### Niche modeling approach

We built an ecological niche modeling pipeline in R to predict the ecological niche of 1,013 species that had at least five occurrence records after undergoing the coordinate cleaning described above. This pipeline adapts the workflow outlined in Abbott et al. (2022) and was designed to automate the building of ecological niche models, while including steps that customize models for each species. First, accessible area, which was the area where the distribution model was fit and projected, was determined by buffering an alpha hull around occurrence records that passed all automated and manual filtering steps. The alpha hull was calculated using the getDynamicAlphaHull function from the R package *rangeBuilder* (Davis Rabosky et al., 2016), and the alpha hull was buffered by the larger value of either 75 km or the 80th percentile distance of each occurrence record to its nearest occurrence record. Next, we fit a Maxent model (Phillips et al., 2006, 2017) with default settings using the *dismo* package in R (Hijmans et al., 2017). Maxent is a machine learning algorithm that fits relationships between occurrence records and background points to predictor layers.

Our initial model included 13 bioclimatic variables from 19 available in WorldClim (Fick & Hijmans, 2017), three soil variables provided by the International Soil Reference and Information Center (Batjes et al., 2017), and two topography layers provided by EarthEnv (Amatulli et al., 2018). Variables were selected from these sources on the basis of biological relevance to plant distributions; collinearity was dealt with at the modeling stage as described below. The soil layers were downloaded at the depths of 0–5 cm, 5–15 cm, and 15–30 cm, and the average value of each cell for these given depths was calculated for use in our models. In total, these predictor variables were bio1, bio2, bio4, bio5, bio6, bio8, bio9, bio12, bio13, bio14, bio15, bio16, bio17, elevation, ruggedness, soil nitrogen, sand, and soil organic carbon. Initial variables had a spatial resolution of approximately 1 km at the equator and were aggregated five-fold to the coarser resolution for model building. We calculated the variance inflation factors (VIF) of our initial global models with all 18 variables; if any predictor variable had a VIF greater than 5, we removed the variable with a VIF greater than 5 that contributed the least to the model given its permutation contribution value. This step was repeated until no variables were retained in the model with a VIF greater than five. These species-specific predictor variables were used in the following step below.

We used the R package *ENMeval* (Muscarella et al. 2014) to evaluate many combinations of Maxent models with different tuning parameters in order to optimize model complexity while maintaining predictability. We fit models for each combination of tuning parameters within range multipliers of 0.5, 1, 2, 3, and 4 and feature classes of “linear”, “linear + quadratic”, “linear + quadratic + hinge + product”, and “linear + quadratic + hinge + product + threshold”. Occurrence and background localities were partitioned into training and testing bins using block partitioning. The model with the lowest AICc value was selected as the top model if it had training and validation area under the curve (AUC) greater than 0.7, while in the few cases where those values were less than 0.7, we selected the model with the highest validation AUC as the top model. Top models were converted to predicted presence/absence maps using the tenth percentile rule, where 90% of locations used for training are converted to predicted presences and the lowest 10% of values are deemed absences. This threshold was chosen because our underlying species occurrence data used in model fitting may still have a small proportion of uncertain or poor-quality records, and thus allowing 10% of presumed presences to be outside predicted suitable habitats helps reduce commission error.

### Species richness, relative phylogenetic diversity, and endemism

The thresholded ecological niche models for all species and the phylogeny described above were imported into Biodiverse v. 3.1 (Laffan et al., 2010). These datasets were used to calculate species richness (SR) and relative phylogenetic diversity (RPD). RPD is the ratio of phylogenetic diversity (PD, measured as the sum of branch lengths connecting the terminal taxa present in each location) on the original phylogenetic tree compared to a phylogeny with the same topology but with a transformation imposed to equalize branch lengths (Mishler et al., 2014). Thus, low RPD represents more shallow branches compared to a tree with unstructured branch lengths, whereas high RPD represents more long branches. We opted to focus on RPD given that raw PD displays strong correlation with SR and that SR allowed us to incorporate many more species given more limited sampling of species-level tips in the phylogeny compared to species distribution models. Mapped raw PD is available as a map in Supplementary Fig. S1.

We also calculated the proportion of species in each grid cell engaging in nodulation. This was performed by matching species to a recent comprehensive genus-level database of nodulating species (Kates et al., 2022). The distribution of nodulation is thought to be fairly well-understood in Fagales (Ardley & Sprent, 2021; Pawlowski & Sprent, 2008), resulting in 100% species-level coverage for all included tips in the phylogeny.

### Randomizations and endemism categorization

We used spatially structured randomizations to determine geographic locations where RPD was significantly higher or lower than expected given a random distribution of species. The randomizations were calculated by randomizing species to grid cells while holding constant the richness of each cell and range size of each species. Values of RPD were then calculated for each iteration of a total of 100, which creates a null distribution for each grid cell. A two-tailed test based on percentiles calculated from the null distribution is used to determine whether observed values are significantly higher or lower (alpha level 0.05 in each direction) compared to null distributions.

We also calculated relative phylogenetic endemism (RPE; Mishler et al., 2014), the ratio between measured phylogenetic endemism (PE) and the PE estimated from a phylogeny with equally distributed branch lengths. We then ran a randomization on RPE as a means to categorize different types of phylogenetic endemism using the Categorical Analysis of Neo-And Paleo-Endemism (CANAPE) approach (Mishler et al., 2014). CANAPE first selects grid cells that are significantly high (one-tailed test, alpha level 0.05) in either the numerator or denominator of RPE and then uses a two-tailed test of the RPE ratio (alpha level 0.025 in each direction) to categorize cells as having a high proportion of neoendemics, a high proportion of paleoendemics, a mixture of both types, or no significant endemism. Neoendemics have higher than expected concentrations of range-restricted short branches, while paleoendemics have higher than expected concentrations of range-restricted long branches. Endemism measures and randomizations were calculated in Biodiverse v. 3.1 (Laffan et al., 2010). A map of raw RPE is available in Supplementary Fig. S2.

### Phylogenetic regionalization

Examining turnover of lineages across space offers the opportunity to test traditional hypotheses of biogeographic regions using quantitative methods (Daru et al., 2017). “Phyloregions” are defined here as clusters of areas of Earth possessing a similar phylogenetic composition of Fagales species, based on community distance metrics. We examined range-weighted phylogenetic turnover by calculating a pairwise distance matrix as a basis for clustering, with the purpose of identifying regions containing similar phylogenetic composition (Laffan et al., 2010). In a range-weighted phylogenetic turnover analysis, values of phylogenetic turnover (e.g., phylobeta) are first measured by comparing the lengths of branches of the overarching tree shared and unshared among pairs of cells. Then the phylobeta values are weighted by the fraction of their geographic range found in that location. We manually selected breaks in the dendrogram that determined well-defined groupings of contiguous sets of colored grid cells. Considering the resulting clusters as phyloregions, we compared these to the most authoritative traditional treatment of woody plant biogeographic regionalization (the 35 floristic regions of Takhtajan et al., 1986) and to a recent similar attempt across the vascular plants (Carta et al., 2022). It is important to point out that, while phyloregionalization and floristic kingdom classifications have similar goals of recognizing floristic assemblages via shared diversity, the latter do not explicitly incorporate phylogenetic relationships. Similarly, because clustering methods rely on distance in terms of community composition rather than geographic distance, they importantly also relax the assumption of a spatially contiguous regionalization that is implicit in floristic kingdoms and other similar classifications (see Results; spatial discontinuity complicates the comparison of two of the regions recovered here).

### Environmental associates of grid cell metrics

We fit a series of models under a model choice paradigm to assess the best environmental predictors of RPD, CANAPE endemism categories, and proportion of grid cell species engaging in nodulation. Environmental data were summarized by eight selected predictors chosen based on the biology of the plants following previous work in the clade (Doby et al., 2022; Siniscalchi et al., 2022): aridity (calculated as the UNEP aridity index following Folk et al., 2020), mean annual temperature (bio1), temperature annual range (bio7; included as a measure of temperature seasonality), annual precipitation (bio12), temperature of the driest quarter (bio17; included as a measure of precipitation seasonality), elevation, and three soil predictors: nitrogen content (chosen to study RNS distribution), soil pH, and soil carbon content (the latter two well-known as constraining plant distributions).

Two classes of models were fit: standard generalized linear models (GLMs) and linear mixed models (LMMs). The logic behind including LMMs was as a natural framework for handling spatial autocorrelation as a random effect and partitioning variance separately from the main predictors, treated as fixed effects. Within each model class, we tested either the full eight-predictor set of environmental variables or a reduced set of five predictors, individually chosen within each model class and response, based on the highest magnitude of normalized coefficients in the full GLM. LMM models additionally fit cell centroid latitude and longitude as random effects. Finally, a LMM model was fit without environment, using only latitude and longitude as random effects, in order to verify the predictive power of environmental data for the response.

To summarize, the model set therefore included five distinct models, GLM-full, GLM-reduced, LMM-full, LMM-reduced, and LMM-no environment, for each of three responses: RPD, CANAPE, and proportion of RNS species. For the continuous RPD and proportion of RNS responses, standard GLM and LMM were used. For the categorical CANAPE response, endemism significance categories were lumped to yield a binary comparison with non-significant cells, and the model family was specified as binary with a logit link function. Model choice used standard AIC (Akaike, 1974) in consideration of the large sample sizes.

### Data availability

Scripts to perform the analyses, as well as raw grid cell products, are available on GitHub (https://github.com/ryanafolk/fagales_phylodiversity).

## RESULTS

### Species richness

Species distribution models covered 1,045 species or 90% of all recognized Fagales species, suggesting robust estimates of species richness patterns. As expected, we recovered the highest species richness of extant Fagales in temperate eastern Asia, with secondary hotspots of species richness in eastern North America, montane Mexico, and Malesia (Fig. 1a). In equatorial regions, species richness is high only in the Malesian biogeographic province (sensu Takhtajan et al., 1986) from Indonesia to Papua New Guinea. Fagalean communities in the Southern Hemisphere are relatively species-poor by contrast, with the highest diversity in southwestern Australia. Finally, although not as high in species richness as the foregoing regions, a further hotspot comprises much of Europe.

**Fig. 1.**
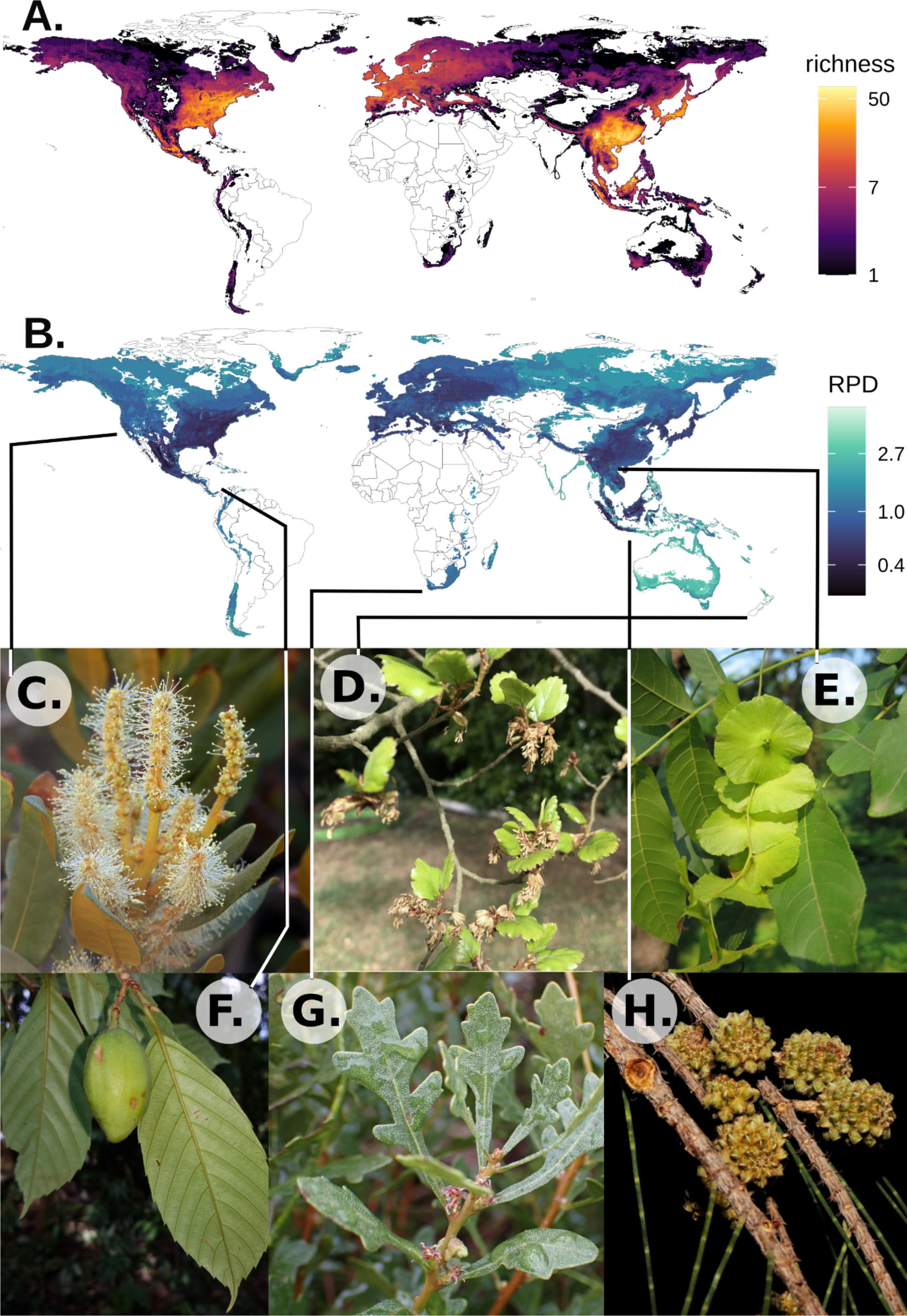
Summary of diversity in Fagales. A. Global species richness of Fagales, with warm colors representing more diverse areas. White areas of land indicate no mapped species. B. Global distribution of relative phylogenetic diversity (RPD) in Fagales, with greener colors indicating more diverse areas. C–G, photos of representative plants: C. *Chrysolepis sempervirens* (photo credit: Steve Matson). D. *Ticodendron incognitum* (credit: Leonardo Álvarez-Alcázar). E. *Nothofagus fusca* (credit: Nicola Baines). E. *Myrica quercifolia* (credit: David Hoare). F. *Cyclocarya paliurus* (credit: Yao Li). G. *Casuarina equisetifolia* (credit: Savvas Zafeiriou).

### Relative phylogenetic diversity

The distribution of RPD was markedly different from that for species richness, showing centers of RPD primarily in the Southern Hemisphere (Fig. 1b). The highest RPD was seen in southern Australia, and relatively high RPD was also associated with eastern and southern Africa, the Andes, and to a lesser extent the boreal Northern Hemisphere. Further minor areas of high RPD, although associated with species-poor communities (Fig. 1a), extend in a narrow area comprising coastal regions in South and Southeast Asia.

### Significance tests of RPD

RPD randomizations demonstrated significant spatial structuring of sites outside null expectations. Areas of significantly high RPD (blue cells, Fig. 2a), corresponding to communities with especially long branch lengths, occur in six biogeographic areas, listed in descending spatial extent: southern Australia, southeast Asia, western Europe, Malesia, the Sierra Madre Oriental south to Central America, and Tierra del Fuego. In the Northern Hemisphere, areas of significantly low RPD (red cells, Fig. 2a) were most prevalent in the Americas, extending from the coastal plain of the southeastern United States to most of montane Mexico. In Eurasia, low RPD was associated with the Mediterranean basin, an additional northern region from Poland to the Volga, and further isolated Asian occurrences mostly in the Himalayan region to montane southeast Asia. In equatorial regions, significantly low RPD was prevalent in the western Malesia biogeographic province (i.e., Sundaland), but significantly low RPD does not occur in the Southern Hemisphere outside of equatorial Malesia.

**Fig. 2.**
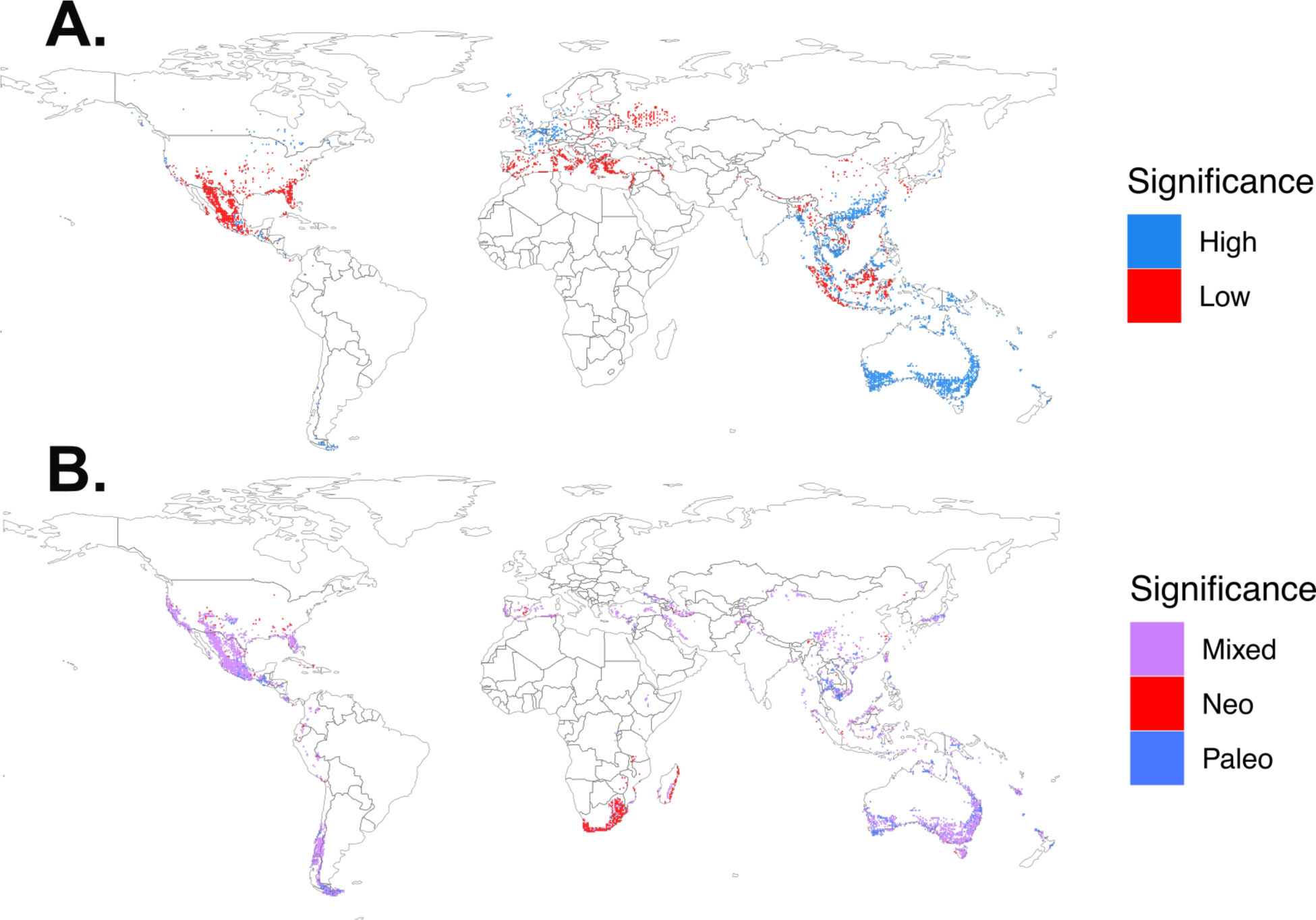
A. Randomization tests for RPD. “High” significance refers to cells above null expectations; “low” significance refers to cells below null expectations. B. CANAPE analysis. Interpretation is similar to (2a) except that CANAPE also distinguishes centers of mixed endemism, which contain species both above and below null expectations.

### Regionalizing neo- and paleoendemism

CANAPE analyses (Fig. 2b) were mostly similar to RPD randomizations (which, although similar in implementation, do not incorporate range size and are not measures of endemism), with CANAPE showing that most significant regions of endemism are characterized by mixed endemism patterns. Southern Mexico, southeast Asia, southwestern Australia, and the southern Andes were characterized by significant paleoendemism. The most important difference between the CANAPE and RPD results was that portions of southern Africa and Madagascar have strong fagalean neoendemism but no RPD significance.

### Phylogenetic regionalization

Fagalean phylogenetic regionalization accorded well with traditionally recognized biogeographic regions as summarized by (Takhtajan et al., 1986), which were rooted primarily in woody plant distributions. The most widely distributed region was a boreal region across the Northern Hemisphere (dark green, Fig. 3), and three more major regions were at mid-higher latitudes in the south boreal region (light green, yellow, dark blue, Fig. 3). The last of these (dark blue), termed here an “Afro-boreal region,” was the only phyloregion present in large areas of both hemispheres, being present in both lower boreal latitudes and in southern Africa, and therefore closely matching areas of Fagales distribution dominated by *Myrica* of Myricaceae.

**Fig. 3.**
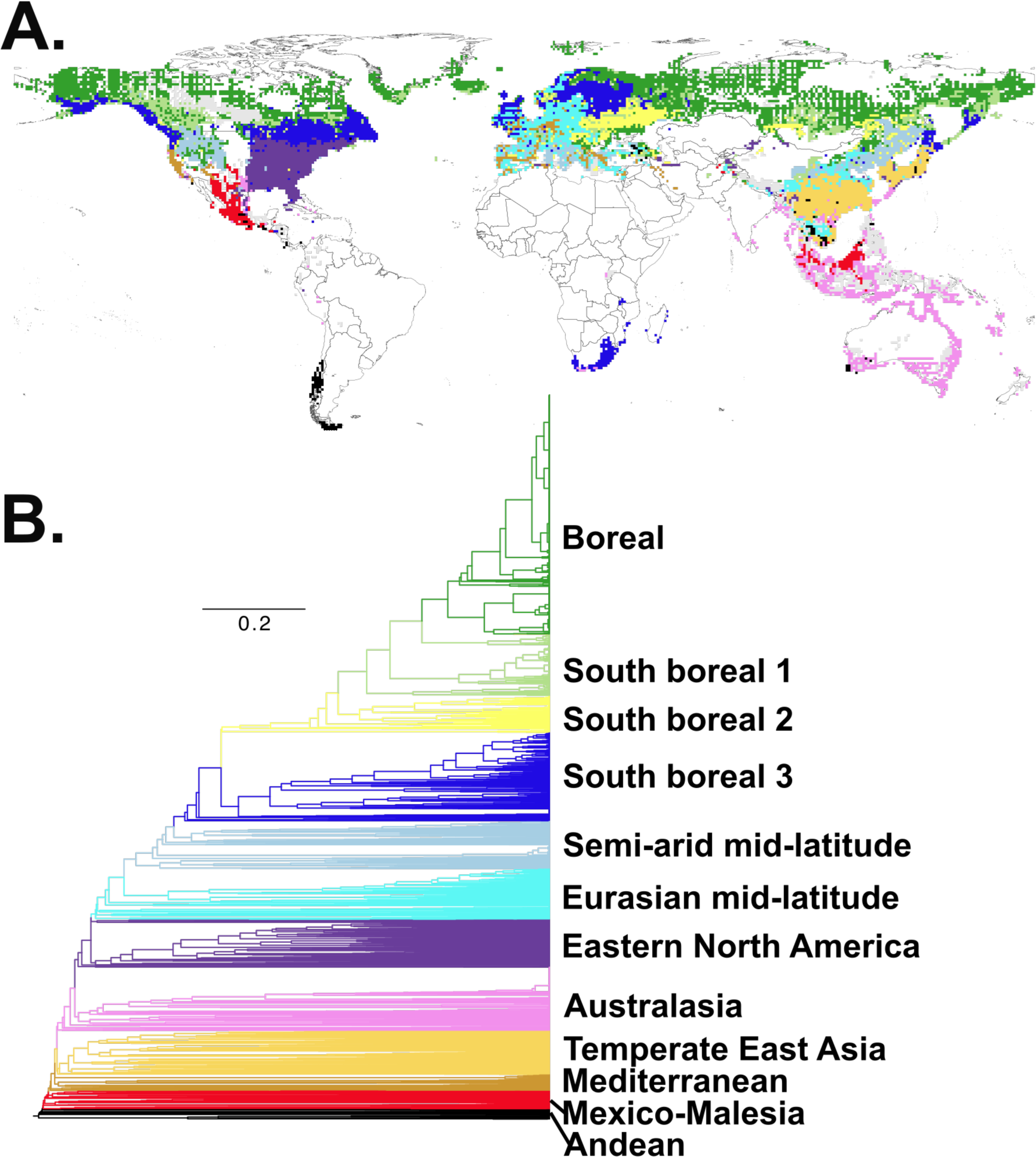
A. Phylogenetic regionalization. B. Corresponding dendrogram showing group colorization and distances. Each terminal represents a grid cell in the map.

Eastern North America and temperate eastern Asia (respectively dark purple and light orange in Fig. 3) were recognized as distinct phyloregions for Fagales, and were not closely related to each other (see dendrogram, Fig. 3b). North America comprises mostly the boreal and eastern North America regions discussed above, but the California Floristic Province was retrieved as part of a distinctive “Mediterranean” phyloregion also occurring in the Mediterranean Basin (Fig. 3, brown), and much of interior western North America was part of a semi-arid mid-latitude region distributed across the Northern Hemisphere (Fig. 3, light blue). Montane Central America was primarily delimited as a phyloregion shared with portions of Malesia (Fig. 2a, red, the second-most divergent phyloregion; Fig. 2b), with a small amount of the eastern North America region in the Sierra Madre Oriental. The semi-arid phyloregion, along with a broadly distributed Eurasian phyloregion (Fig. 3, turquoise) comprised continental Europe and the Mediterranean Basin.

Two phyloregions were unique to the Southern Hemisphere. One is a widespread region of Malesia and coastal Australia east to northern New Zealand (Fig. 3, pink), representing Fagales communities dominated by Casuarinaceae. The second is a phyloregion unique to southern South America (Fig. 3, black) that reflects the distribution of Nothofagaceae in Valdivian forests; this was the most distinctive phyloregion (Fig. 3b).

### Distribution of RNS

The distribution of species of Fagales engaged in RNS (Fig. 4) is highly heterogeneous and nodulating species (those displaying the characteristic root structures of RNS) distributed in areas of low Fagales species richness but associated with areas of high RPD (Figs. 1-2). A particularly high richness of Fagalean nodulators (but not overall Fagalean species richness) exists in southern Africa (reflecting a local radiation of Myricaceae) and the southern Malesian region and Australia (reflecting the primary distribution of Casuarinaceae); in these primarily semi-arid habitats, no non-nodulating Fagales are known. A secondary area of high nodulator proportions occurs across boreal latitudes in the Northern Hemisphere, with lower nodulating species diversity than the Southern Hemisphere but more site-level co-occurrence of distinctive lineages from both Betulaceae and Myricaceae.

**Fig. 4.**
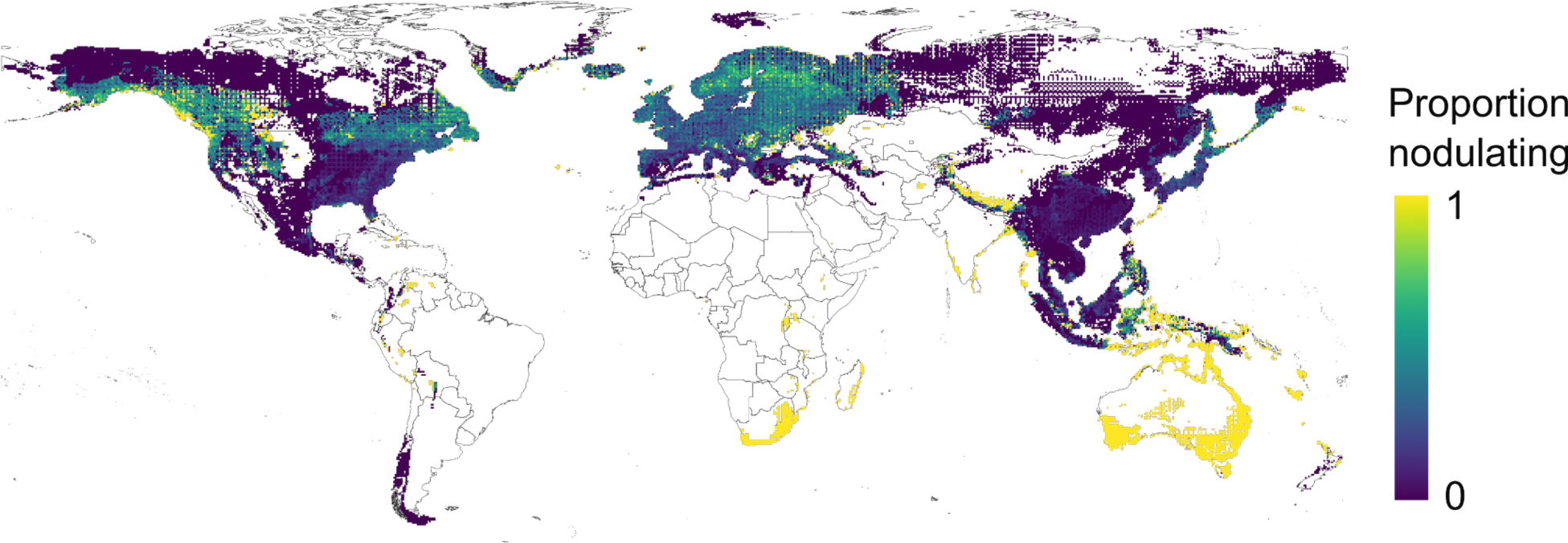
Proportion of grid cell species engaging in RNS (root nodule symbiosis). Note that areas with high nodulator proportions correspond with areas of high RPD (Fig. 1b).

### Environmental associates

For RPD, CANAPE, and proportion of nodulators, model selection via AIC favored the full mixed model with latitude and longitude. Aridity was the most important predictor for all responses.

The favored full mixed model for RPD (ΔAIC = 14.7669; Supplemental Table S1) had strong explanatory power (conditional *R*^2^ = 0.8656), but the fixed environmental effects only had weak explanatory power for RPD (marginal *R*^2^ = 0.0211). Nevertheless, models with environmental predictors are strongly favored (best model vs. no-environment model, ΔAIC = 619.859). Comparing among the predictors with normalized coefficients, the most important predictor was aridity (positive relationship, meaning higher RPD in more mesic sites), followed by soil carbon and pH (Supplemental Table S2).

Similarly, environmental factors had a moderate amount of explanatory power for CANAPE significance categories (marginal *R*^2^ = 0.3007; conditional *R*^2^ = 0.6388), where the complex mixed model was favored (ΔAIC = 31.43719). Among these, arid environments were consistently associated with all forms of endemism significance (Fig. 6a; significant endemism associated with more arid sites). Similarly to RPD, models with environmental predictors are strongly favored (best model vs. no-environment model, ΔAIC = 276.6233; Supplemental Table S1).

The proportion of the community engaging in nodulation had similar explanatory power to RPD (marginal *R*^2^ = 0.0467; conditional *R*^2^ = 0.8063) with the full mixed model favored (ΔAIC = 1602.14). Aridity was the most important predictor (Supplemental Table S2), with more mesic environments associated with more nodulators (Fig. 5d). This result seemed to conflict with high-nodulator sites in the Southern Hemisphere (Fig. 4), so the analysis was rerun as two separate models partitioning sites above and below the equator. This confirmed that the aridity relationship differed by hemisphere, with the Northern Hemisphere showing a positive relationship (univariate *R*^2^ 0.03804, *p* < 2.2e-16; i.e., more nodulators in more mesic environments) and the South Hemisphere negative (univariate *R*^2^ 0.3885, *p* < 2.2e-16; i.e., more nodulators in more arid environments; see Fig. 5d).

**Fig. 5.**
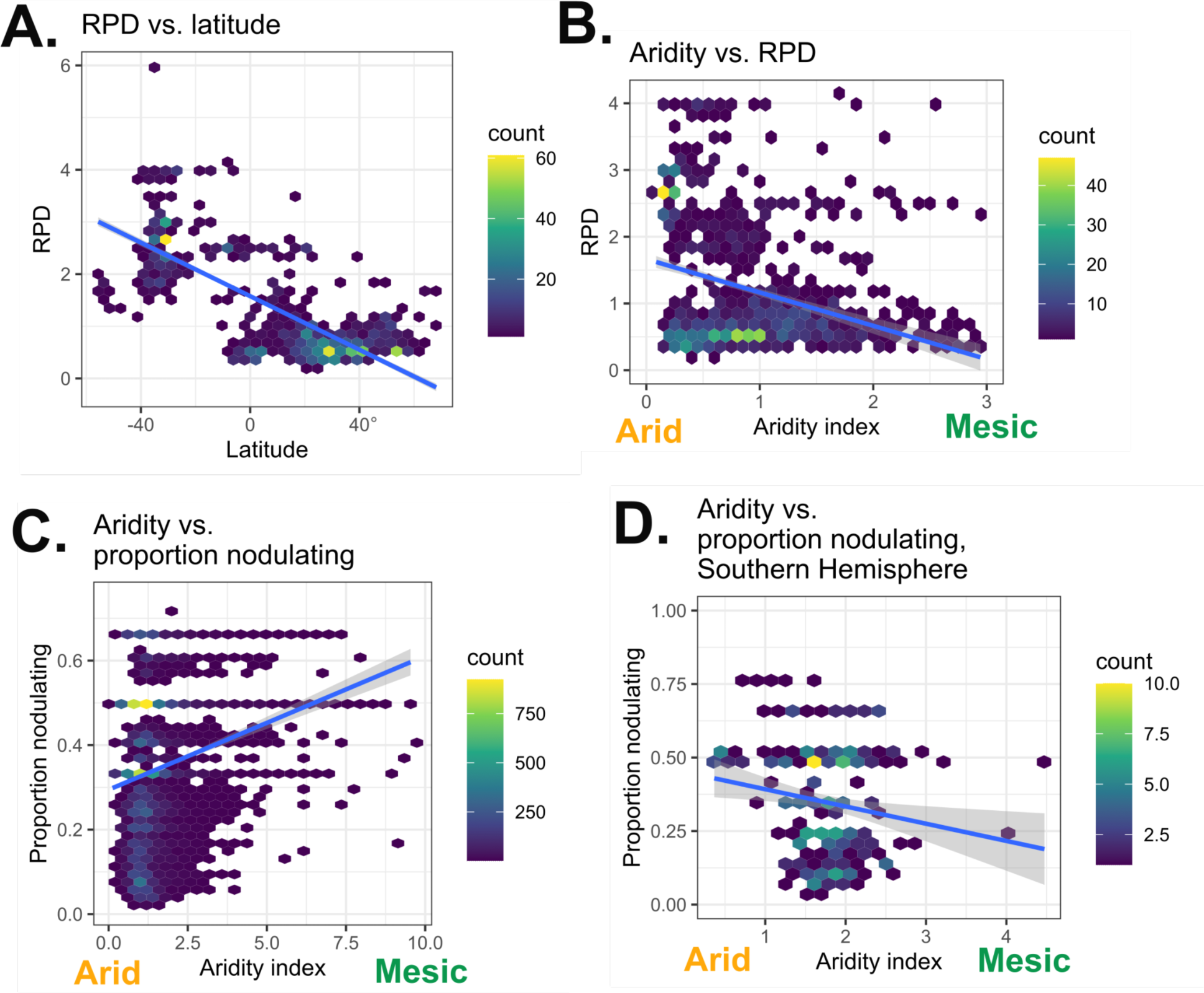
Environmental associates of richness and RPD. Hex points in scatter plots represent aggregations of individual grid cells, with cell density as indicated by the color scale. A. RPD vs. latitude; notice the monotonic behavior of the response and the existence of two clear RPD clusters by latitude. B. Aridity vs. RPD. C. Aridity vs. proportion nodulating; inflated 0 and 1 values omitted per Methods. D. Aridity vs. proportion nodulating, as in (C) but showing only the Southern Hemisphere.

**Fig. 6.**
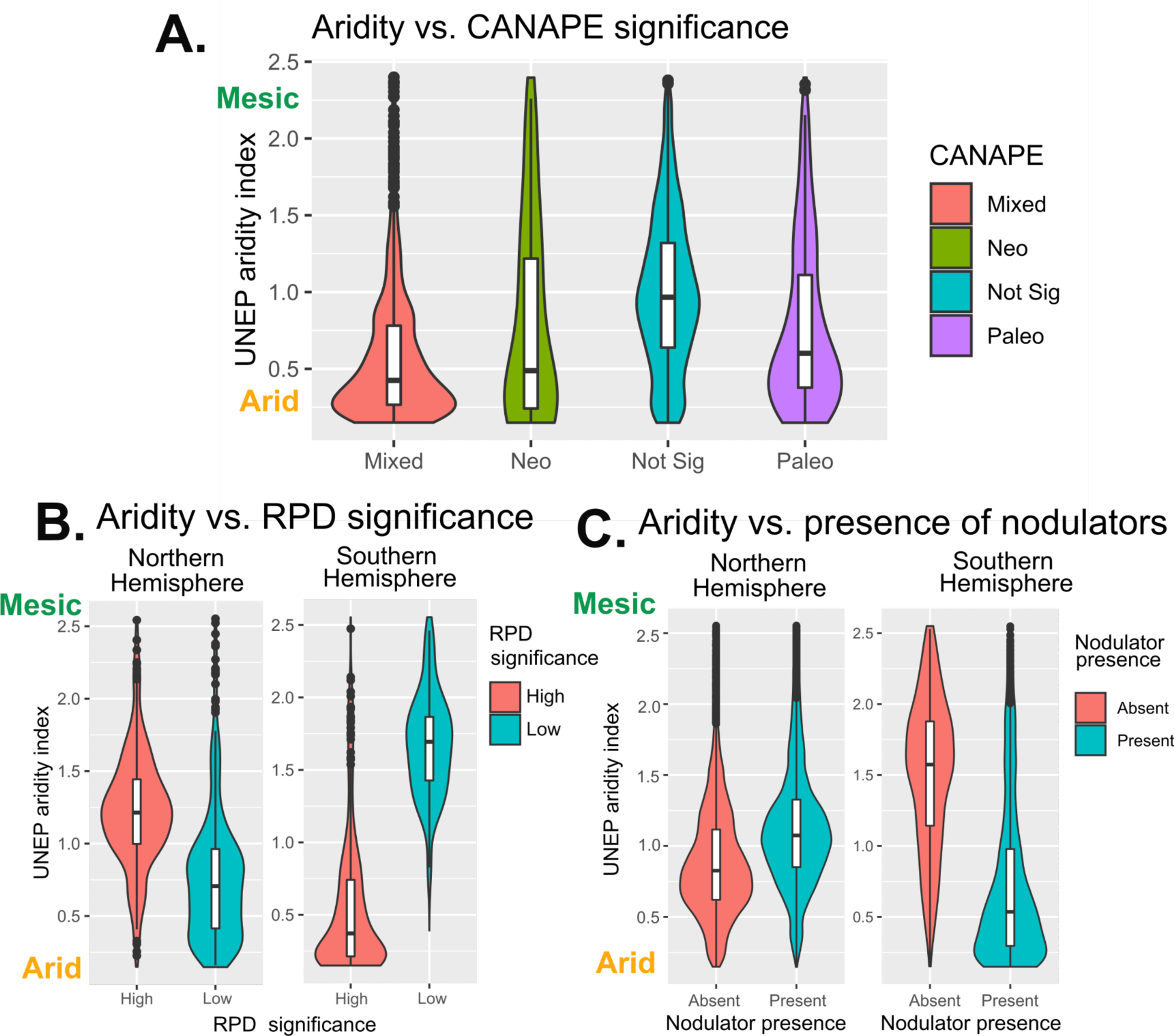
A. Aridity vs. CANAPE significance categories. B-C: breakdown of aridity relationships between the Northern (left) and Southern (right) Hemispheres. B: Low RPD significance (branch lengths significantly less than random expectation) vs. aridity. C: Site-based presence/absence of nodulators vs. aridity.

## DISCUSSION

### Distribution of Fagalean diversity

A major finding of this study was that while centers of species richness for Fagales are similar to those of other Northern Hemisphere temperate plants, relative phylogenetic diversity peaks in low SR regions of Fagales distributions in the Southern Hemisphere. These RPD results, with some of the most species-poor areas being highest in RPD, are unanticipated. High RPD was most spatially extensive at relatively high latitudes (Fig. 1b). However, Australia and adjacent areas stood out as high-RPD outliers while boreal regions were within null expectations (Fig. 2b). Significantly high Southern Hemisphere RPD therefore drives a surprising relationship with latitude, increasing monotonically southwards (Fig. 5a), an unusual result since in many clades, the Southern Hemisphere is lower in diversity (Economo et al., 2018). Accordingly, most Southern Hemisphere cells with significantly high RPD represent distinct lineages of only one family, Casuarinaceae. This result could be interpreted as evidence for ecological filtering as southern Africa and Australia (but not South America) primarily consist of nodulating species (below) and sites with significantly high RPD show a close correspondence with the distribution of nodulation.

### Phylogenetic regionalization

Our results are partly consistent with regionalizations previously identified in other clades, identifying Northern-Southern Hemisphere distinctions as a fundamental divide (Carta et al., 2022). The main finding of Carta et al. (2022), an analysis across vascular plants, was that phylogenetic clustering partly supports traditional biogeographic provinces but solely recognizes a Northern-Southern Hemisphere divide. We found similar distinctive communities in each hemisphere for the Americas and Australasia-Malesia, but we find that the Fagalean flora of southern Africa is most similar to floras in boreal regions, together forming an Afro-Boreal flora. Likewise, the Asia/Pacific divide between hemispheres is different in this study as Malesia and Australasia are not recovered as distinct provinces. This southern rather than equatorial affinity of Malesia may be a special feature of Fagalean biogeography as it is not generally recovered in vascular-plant-wide studies (Carta et al., 2022), although a partly similar regionalization was recovered in mosses (Sanbonmatsu & Spalink, 2022).

In contrast to traditional concepts of temperate diversity, the temperate East Asia and eastern North America phyloregions are not closely related and form separate, strongly distinct floras (see Fig. 3b dendrogram). The boundaries of these two phyloregions accords well with their traditional definitions; eastern North America is almost identical to Takhtajan’s North American Atlantic region (Takhtajan et al., 1986). Our east Asia region is similar to but narrower than Takhtajan’s Eastern Asiatic region, including Japan and the Korean peninsula but excluding northeastern China, Sakhalin, and the Ryukyus, and including only portions of central Hokkaido. This north-south delimitation is very similar to a recent phyloregionalization in China (Ye et al., 2019) but there was no sign of a major divide between eastern and western China (cf. Lu et al., 2018; Ye et al., 2019).

As an additional point, Carta et al. (2022) interpreted distinctive Northern vs. Southern Hemisphere floras as reflective of Laurasian vs. Gondwanan floras. While Fagales is old enough as a clade to be consistent with this interpretation (Larson-Johnson, 2016; Siniscalchi et al., 2023), none of the geological records of subclades of Fagales are completely consistent with this. Casuarinaceae and Nothofagaceae are controversial and may both be post-Cretaceous radiations (Larson-Johnson, 2016; Siniscalchi et al., 2023 but see Sauquet et al., 2012; Wheeler et al., 2022); the distribution Myricaceae in southern Africa and the boreal region also likely reflects a post-Cretaceous radiation (Larson-Johnson, 2016; Siniscalchi et al., 2023). Overall, north and south distinctions are important, but which geological processes enforce this, or whether it arises from fundamental physiological constraints or other forces (Folk et al., 2020; Zanne et al., 2018), requires more research.

The main distinction of species richness in Fagales compared to traditionally recognized centers of temperate forest diversity is the high diversity of montane equatorial areas in Central America and Malesia, more usually considered relictual and marginal areas of (climatically) temperate diversity (Miranda & Sharp, 1950). This finding accords, however, with the distribution of many isolated lineages in Fagales treated as species-poor or monotypic taxa. Montane Central American flora and much the Malesia region were recovered as a single phyloregion, reflecting the shared fagalean diversity among these distant regions. The remainder of Malesia is primarily part of an Australasian region, reflecting Casuarinaceae and other taxa primarily distributed in the southern hemisphere and reaching their northern limit in this area. Mainland Southeast Asia is similar, with a strong north-Asian component and some influence from temperate East Asia, but also is a mosaic of other phyloregions including (most prominently) Australasian influence.

Western and central Europe formed an additional hotspot of species richness that could represent sampling intensity bias. However, high species richness in this area has been identified before in Fagales (Lyu et al., 2022) and is also reflected in a recent investigation of actinorhizal RNS species including Fagales (Tamme et al., 2021: Fig. 2c). Species richness patterns recovered here overall compare closely to those reported in Fagales in a recent methodological contribution (Lyu et al., 2022).

### Actinorhizal nodulators

The relationship of RNS with environment in Fagales (Fig. 1C) was unexpected. Previous studies have supported aridity as favoring RNS (Pellegrini et al., 2016; Siniscalchi et al., 2022) and especially the phylogenetic diversity of RNS (Doby et al., 2022). Whereas most previous studies have treated all nodulators equally and therefore primarily reflect the distribution of the more diverse and prevalent legumes (which are particularly successful in semi-arid habitats), this study represents the first in-depth look at the spatial distribution of phylogenetic diversity specifically for actinorhizal RNS species. The direction of the relationship with aridity found here for Fagales was opposite previous results, with wetter environments showing greater prevalence of RNS species. This result makes sense in light of the specific strategies of nodulators in Fagales that differ from other clades that nodulate. The Southern Hemisphere RNS species of Fagales (Casuarinaceae and Myricaceae) specialize in semi-arid habitats and, although not found the most arid areas, otherwise fit the spatial pattern seen in legumes. However, there is more lineage diversity and a more extensive spatial distribution of fagalean nodulators in the Northern Hemisphere (Fig. 4), and the species involved (Myricaceae and Betulaceae) are often semi-aquatic.

Segregating the results by hemisphere confirms that RNS species have differing ecological strategies within Fagales, with more nodulators in more mesic environments in the Northern Hemisphere and more nodulators in more arid environments in the Southern Hemisphere. These modeling results focused on raw RPD but are robust to randomizations: significantly high RPD is strongly associated with more arid environments in the Southern Hemisphere, but with more mesic environments in the Northern Hemisphere (Fig. 6b). Similarly, when contrasting sites that either possess or lack RNS plants, arid sites are overrepresented in the Southern Hemisphere but underrepresented in the Northern Hemisphere (Fig. 6c). These results suggest that arid specialization is not universal in RNS species and is conditional on local radiations in different biogeographic provinces with differing ecological contexts and bacterial partnering. These results accord with already recognized ecological differences between rhizobial and actinorhizal RNS species (Ardley & Sprent, 2021; Folk et al., 2020; Siniscalchi et al., 2022).

### Conclusions

We found support for traditionally recognized regionalizations for north temperate forests (east Asia, eastern North America), but investigations of RPD and endemism identify novel hotspots including Malesia and central and southern Mexico, not traditionally considered centers of ancient diversity for temperate groups. Phylogenetic regionalization also corresponded to traditional concepts, but some phyloregions were unexpectedly distinct. Potential drivers of nodulating plant distributions disagree with previous work and suggest differing ecological strategies between Northern and Southern Hemisphere species in Fagales. Overall, our results point to the importance of clade-level investigations of phylogenetic diversity for investigating evolutionary processes.

### Contributions

RAF, RPG, and MB conceived the study in conversation with all authors. MB performed occurrence record gathering and cleaning, species distribution modeling, randomization and clustering analyses. CMS performed phylogenetic analysis. RAF performed site-level statistical modeling and wrote the first draft with MB. All authors contributed to the final draft.

