## Supplement for "Spatial phylogenetics of Fagales: Investigating the history of temperate forests"

**Table S1.** Model choice.

| Model | RPD AIC | CANAPE AIC | Proportion nodulator AIC |
|---|---|---|---|
| GLM, full model | 2993.643 | 1270.424 | 43024.94 |
| GLM, simple model | 2992.009 | 1296.337 | 45469.65 |
| LMM, full model | **1688.108** | **1168.315** | **29419.48** |
| LMM, simple model | 1711.882 | 1199.752 | 31021.62 |
| LMM, no-environment model | 2307.967 | 1444.938 | 44315.39 |
| ΔAIC | 23.774 | 31.437 | 1602.14 |

**Table S2.** Model parameters.

| **Model predictor** | **Best RPD model normalized coefficient** | **Best CANAPE model normalized coefficient** | **Best proportion nodulation normalized coefficient** |
|---|---|---|---|
| UNEP aridity index | **0.13026** | **2.70875** | **0.178740** |
| Bioclim 1 | 0.07517 | 0.45883 | -0.117995 |
| Bioclim 12 | 0.02473 | -1.38495 | -0.110906 |
| Bioclim 7 | 0.01945 | 1.21440 | -0.043118 |
| Bioclim 17 | -0.09180 | 0.45492 | -0.090575 |
| Nitrogen content | -0.04362 | -0.31028 | -0.048296 |
| pH | 0.09885 | -0.05209 | 0.175306 |
| Organic carbon content | 0.12994 | 0.14216 | 0.086012 |

**Fig. S1.** Global Fagales PD (phylogenetic diversity).

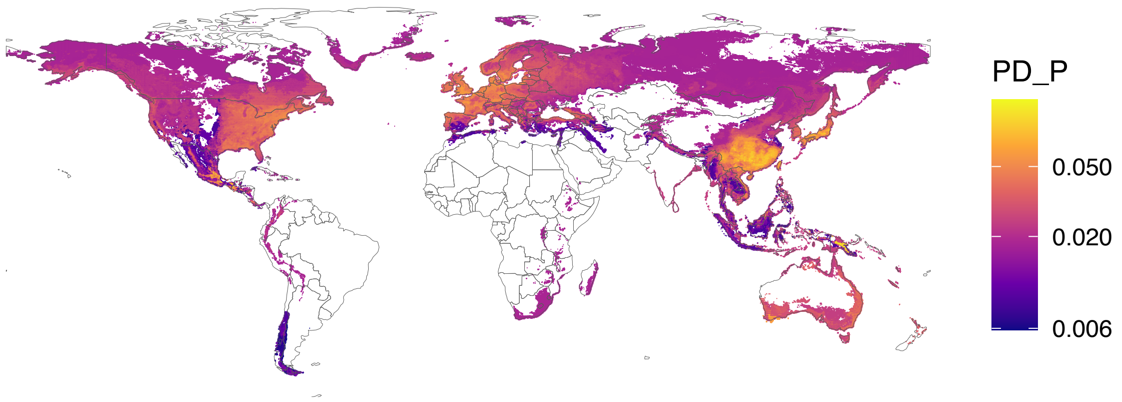

**Fig. S2.** Global Fagales RPE (relative phylogenetic endemism).

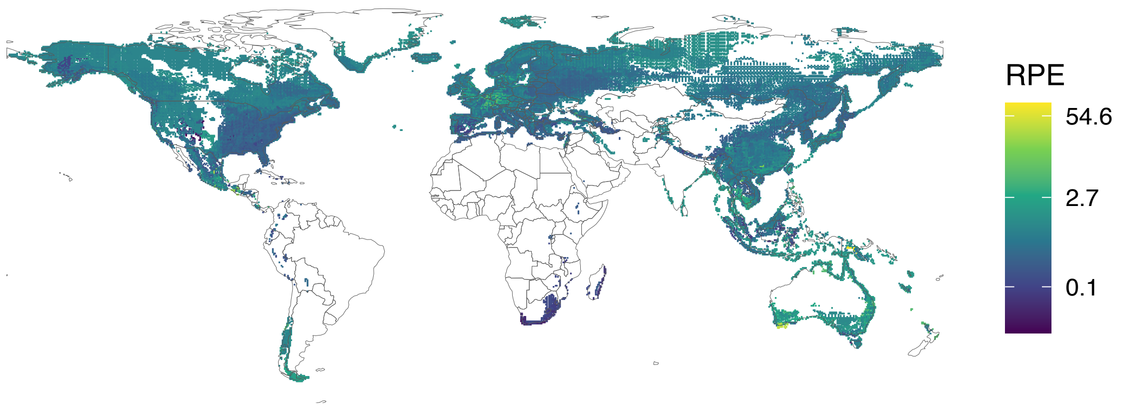
